# Optogenetic stimulation recruits cortical neurons in a morphology-dependent manner

**DOI:** 10.1101/2024.03.18.585466

**Authors:** David Berling, Luca Baroni, Antoine Chaffiol, Gregory Gauvain, Serge Picaud, Ján Antolík

## Abstract

Single-photon optogenetic stimulation is a crucial tool in neuroscience, enabling precise, cell-type-specific modulation of neuronal circuits. Miniaturization of this technique in the form of fully implantable wide-field stimulator arrays enables interrogation of cortical circuits in long-term experiments and promises to enhance Brain-Machine Interfaces for restoring sensory and motor functions. However, for both basic science and clinical applications, it is essential that this technique achieves the precision needed for selective activation of sensory and motor representations at the single-column level. Yet studies report differing and sometimes conflicting neuronal responses within the stimulated cortical areas. While recurrent network mechanisms contribute to complex responses, here we demonstrate that complexity starts already at the level of neuronal morphology. Simulating optogenetic responses in detailed models of layer-2/3 and layer-5 pyramidal neurons, we accounted for realistic physiological dynamics across different stimulation intensities, including threshold, sustained, and depolarization-block responses. Our findings suggest that the spatial distribution of activated neurons from a single stimulator location at the cortical surface can be inhomogeneous and varies with stimulation intensity and neuronal morphology across layers, potentially explaining the observed response heterogeneity in earlier experiments. We found that activation spreads laterally up to several hundred micrometers from the light source due to neuronal morphology. To enhance precision, we explored two strategies: preferentially somatic expression of channelrhodopsin, which was effective only in layer-5 neurons, and narrowing the stimulating light beam, which improved precision in both layers. Our results indicate that, under the right optical setup, single-column precision of stimulation is achievable, and that optical enhancements to the stimulator may offer more significant precision improvements than genetic modifications targeting the soma.

## Introduction

Optogenetic stimulation offers neuroscientists a unique tool for manipulating neuronal activity with precision, providing cell-type specificity and high spatial and temporal resolution (1, 2). Until recently, long-term optogenetic wide-field manipulations in higher mammals were limited, but now become feasible through implantable stimulator arrays using single-photon-based stimulation (3, 4). This advancement opens a path towards a sophisticated interrogation of sensory or motor code in meaningful volumes of cortex (3, 5–8), as well as development of neuro-prosthetic systems for vision and hearing loss remediation (3, 8–10). However, these applications require spatially precise stimulation to selectively engage functional cortical representations at a spatial scale of cortical columns (11, 12) requiring sub-millimeter precision (13), which in experiments proved challenging due to the spatial spread of stimulation-evoked responses (5, 6, 8). The observed spatial spreading and complex distribution of responses to stimulation may be attributed to propagation and transformation of the stimulation-evoked responses in the network (14–17). Alternatively, studies on neuron excitability suggest that optogenetic activation can recruit neurons via distal parts of their morphology (18, 19), raising the question whether direct activation of distant neurons via their spatially extending morphology additionally drives spread-out and spatially complex responses to optogenetic stimulation?

Here, we approached this question using computational methods to reveal how optogenetic responses of a layer-2/3 and a layer-5 pyramidal neuron depend on the lateral displacement of an optical fiber placed on the cortical surface. We validated the model’s basic optogenetic response dynamics against data from primate retinal ganglion cells (RGCs) recorded across a broad range of stimulation intensities, inducing threshold, sustained and depolarization block responses.

Our simulations demonstrate that since distant parts of neuron morphology contribute to activation, optogenetic stimulation recruits neurons distributed in a complex manner in the cortical network beneath the stimulator. We further find that non-linearities in the optogenetic mechanism and neuronal response function cause a stimulation-intensity-dependent redistribution of the activated neurons. Further, we find that due to neuronal morphology, different spatial patterns of cells will be optogenetically activated in layer-2/3 than in layer-5. These findings reveal that neuronal morphology causes a diverse stimulation-intensity and layer-dependent spatial distribution of optogenetic responses, that can explain the experimentally observed variation of stimulation outcomes.

The simulations predict that surface optogenetic stimulation significantly activates layer-2/3 neurons in 100 µm and layer-5 neurons in 200 µm distance to the stimulator, which suggests that surface stimulation may enable selectively recruiting functional neuronal representations encoded at a submillimeter scale of cortical columns. Yet, higher precision may be necessary, since evoked activity could spread beyond the directly activated cortical volume through synaptic transmission in the network. We therefore investigated whether genetics-based or optical improvements to the optogenetic setup could further enhance precision. We simulated soma-targeted channelrhodopsin expression derived from fluorescence measurements in cortical brain slices, finding that spatial precision was only improved for the layer-5 neuron and high channelrhodopsin expression. On the other hand, increasing the collimation of the optical setup improved spatial precision irrespective of the neuron-type and maximized spatial stimulation precision for the tightest light beam explored regardless of whether channelrhodopsin targeting to cell bodies was involved. These results imply that optical improvements of the stimulator are more effective in increasing precision of surface-based stimulation than genetics-based targeting of the opsin to the cell body.

## Results

### Computational Model

We simulated optogenetic neuronal responses of channelrhodopsin-2 (ChR2) expressing pyramidal neurons from cat primary visual cortex layers 2/3 and 5. Neuronal electrophysiology was simulated with the NEURON simulator (20) based on previously developed multi-compartment neuron models (18, 21, 22) [Fig. 1A,B]. Our model further uses an optical fiber light model to calculate light intensities along the neuronal membrane accounting for attenuation through scattering and absorption (23) [Fig. 1A] as well as a model of channelrhodopsin light activation (24) to predict optogenetically induced membrane conductance [Fig. 1C]. We approximated the distribution of channelrhodopsin throughout the cell morphology as uniform in agreement with experimental data (25, 26). Key variables of the simulation workflow are illustrated in Fig. 1D. Details of the model are described in Methods.

**Figure 1:**
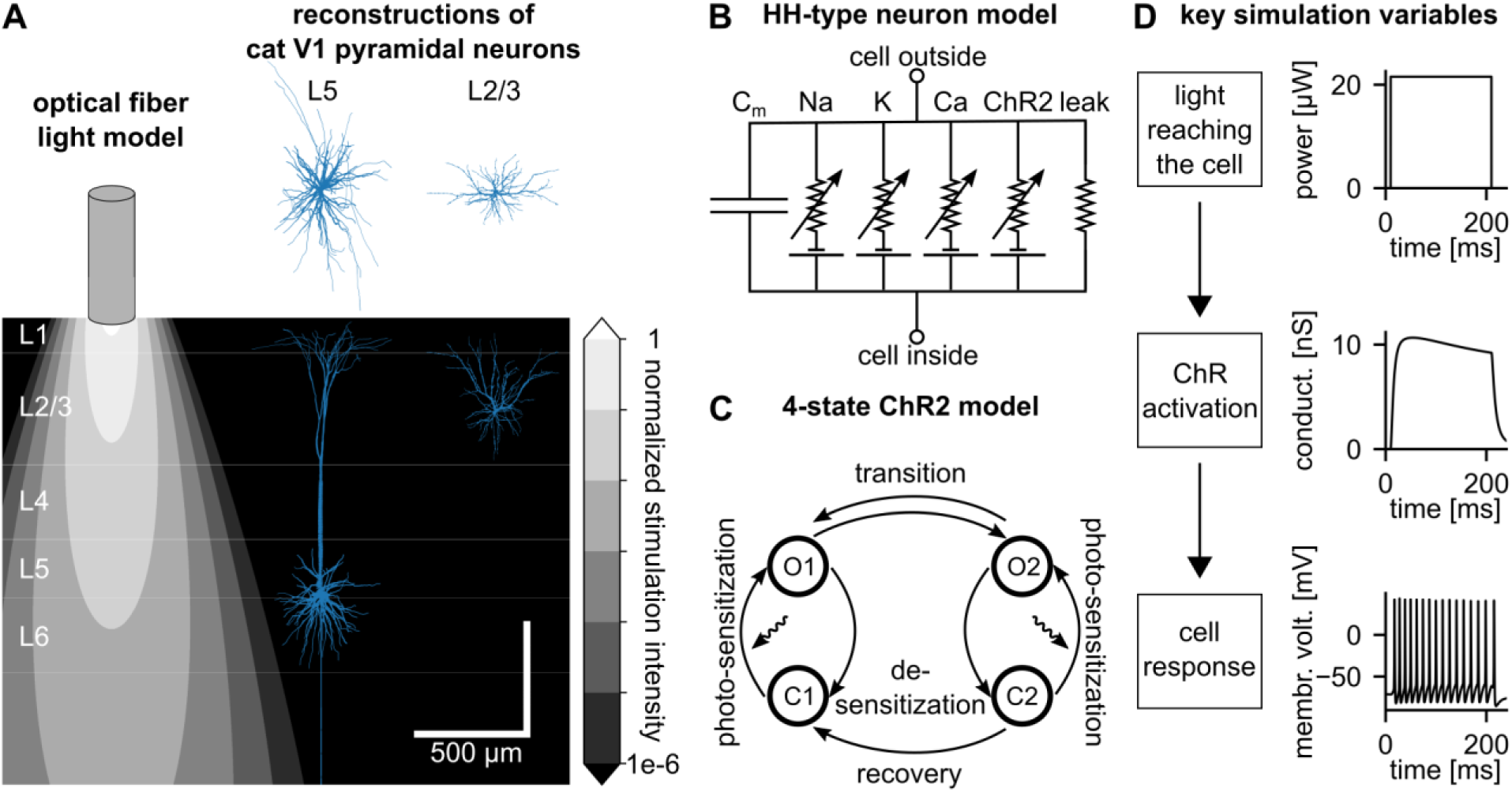
Optogenetic response model. **(A)** Simulated light intensity distribution of an optical fiber (18, 23) and reconstructions of the layer-5 and layer-2/3 pyramidal neuron morphology (18, 21) in proportion to the cortical volume with layers indicated on the left. **(B)** Hodgkin-Huxley-type neuron model with membrane mechanisms used in this study (18, 22). **(C)** 4-state Markov model used to simulate the activation of Channelrhodopsin-2 (ChR2) (18, 24). **(D)** Behavior of key variables during an example stimulation of the layer-5 neuron: Light power reaching the cell membrane results in conductance through ChR2 channels, and finally causes a neuronal response. Plots show sums across the cell membrane for light power and ChR2 conductance as well as the membrane voltage at soma for the cell response. Stimulation intensity was 9.5 mW/mm² measured at stimulator output surface.

### Model optogenetic responses match cell-attached recordings of optogenetic responses in primate RGCs

We verified that our model dynamics of optogenetic responses are biologically plausible by comparing the optogenetic response of the simulated layer-5 neuron to cell-attached recordings of transduced primate retinal ganglion cells (RGCs) [Fig. 2]. Optogenetic responses of RGCs were recorded for absolute stimulation intensities ranging between 4e14 photons/cm²/s to 3e17 photons/cm²/s measured at the neuron’s soma. To account for variations in opsin expression, efficiency, and light attenuation, we compared stimulation intensity levels on a normalized scale. Biological responses were recorded at the multiplications of 1, 50, 200, 250, and 750 of the lowest stimulation intensity. This intensity range induced optogenetic responses covering threshold response at lowest, sustained response at 50-fold, and depolarization block at 200-750-fold normalized stimulation intensity. This behavior was observed in about half of the cells recorded from one retina. Remaining cells had varied responses but did not exhibit depolarization block within the recorded range of stimulation intensities, which may have been caused by lower opsin expression, and therefore insufficient stimulation intensity to reach depolarization block [see Supplementary Materials Fig. S4]. Model responses simulated at an expression density of 130 channels/µm² and the same (normalized) stimulation intensity steps captured response onset, spike count, and amplitude modulation of the biological responses of cells exhibiting depolarization block. For example, RGC 1 responded with a single spike at about 30 ms latency at lowest and a sustained response of 35 spikes at 50-fold stimulation intensity [Fig. 2A]. At 250- and 750-fold normalized intensity, the cell exhibited few spikes of decaying amplitude within 10-20 ms after stimulation onset. Model responses qualitatively matched the latency of the single spike response (approx. 20 ms) at lowest intensity, the spike count at sustained response (33 spikes) appearing at 50-fold intensity, and a complete decay of spike amplitude within 10-20 ms in the depolarization block response at 250-, and 750-fold intensity [Fig. 2B]. Model responses also agreed with optogenetic responses recorded in another RGC (RGC 2), and further captured the modulation of spike amplitudes occurring during sustained response at 50-fold normalized stimulation intensity [Fig. 2B, bottom]. In summary our model exhibits intensity-dependent changes in response dynamics, including depolarization block, which we validated based on matching observations in a major number of neurons recorded for a range of stimulation intensities from a single Macaque retina.

**Figure 2:**
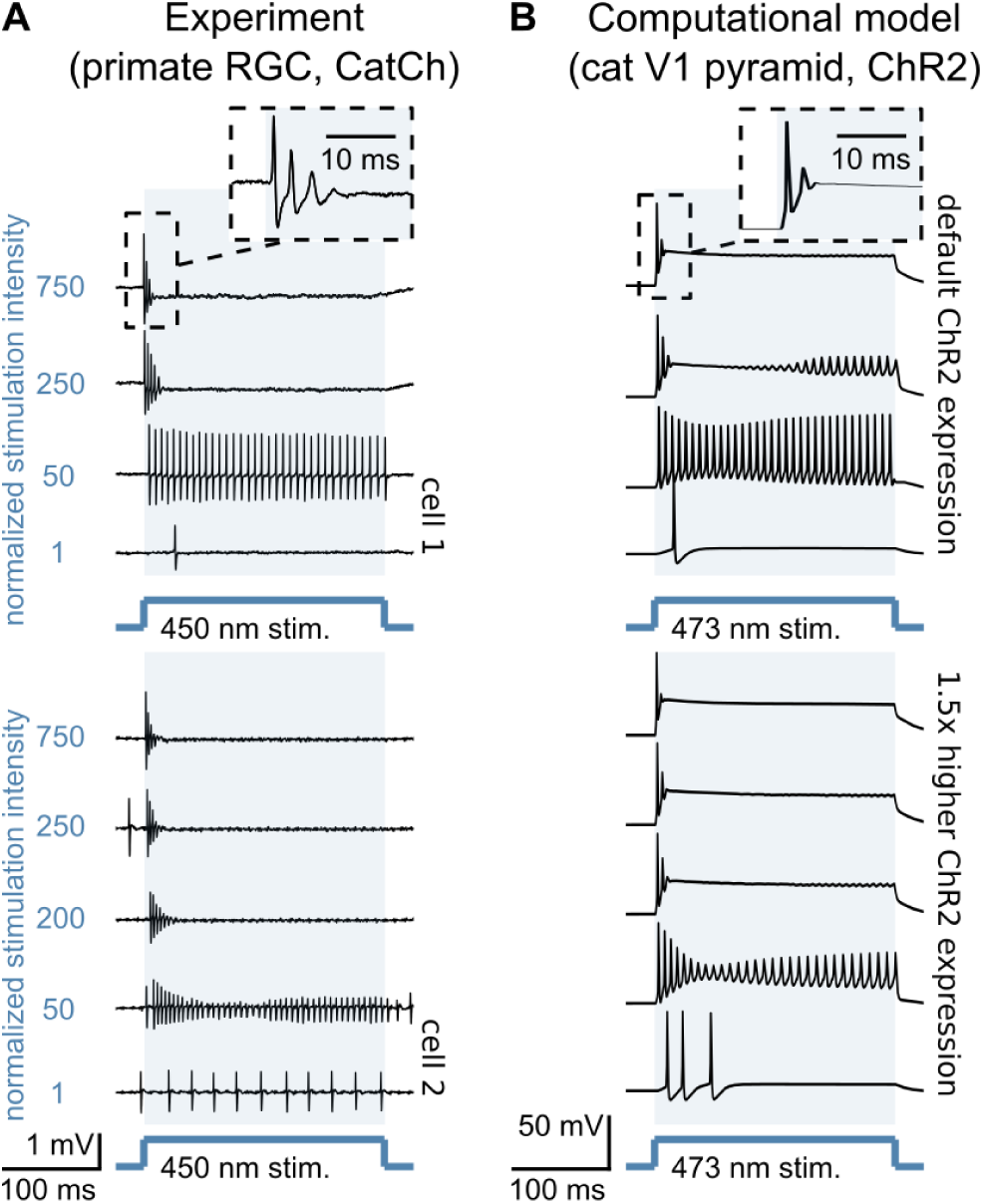
Agreement of model optogenetic responses with biological data. **(A)** Cell-attached recordings of two primate retinal ganglion cells (RGCs) undergoing optogenetic stimulation at stepwise increasing stimulation intensity starting from 4e14 photons/cm²/s (corresponding to a normalized stimulation intensity of 1). **(B)** Model optogenetic responses of the layer-5 pyramidal neuron reconstructed from cat primary visual cortex. Model responses matched responses of RGC cell 1 at a ChR-2 expression of 130 channels/µm² and RGC cell 2 for a 1.5-fold higher expression of 195 channels/µm² across the same range of normalized intensities. To account for differences between the experimental and simulated computational setups, absolute intensities were increased by a factor of 14.75 for the computationally modelled responses.

### Morphology causes heterogenous spatial distribution of optogenetic activation beneath the light source

Keeping stimulation intensity constant, we placed the stimulator at various positions along the cortical surface [Fig. 3A] and simulated optogenetic responses of the layer-5 neuron to a 200-ms duration constant light pulse [Fig. 3B, top]. We counted spikes in somatic membrane voltage through thresholding at 0 mV and quantified firing rate by dividing by stimulation duration. By mapping the optogenetic responses with respect to the position of the stimulator relative to the cell body projected onto the cortical surface [Fig. 3B, bottom], we found that optogenetic responses varied in a heterogenous way with stimulator position. At a stimulation intensity of 16 mW/mm² at the stimulator surface, the neuron’s response was weaker for a central stimulator location above the soma compared to a 200 µm lateral displacement. While exhibiting a decaying trend towards more lateral positions from center, isolated stimulator locations at more than 300 µm distance from the neuron’s soma caused responses of similar strength compared to central positions. These results highlight that spatially extending neuronal morphology of layer-5 neurons could contribute to around 500 µm spread of optogenetic activity along the cortical surface, considering the average full-width at half-maximum of the optogenetic response. Further, individual and spatially isolated responses may introduce noise into spatial stimulation patterns at a larger scale of up to one millimeter.

**Figure 3:**
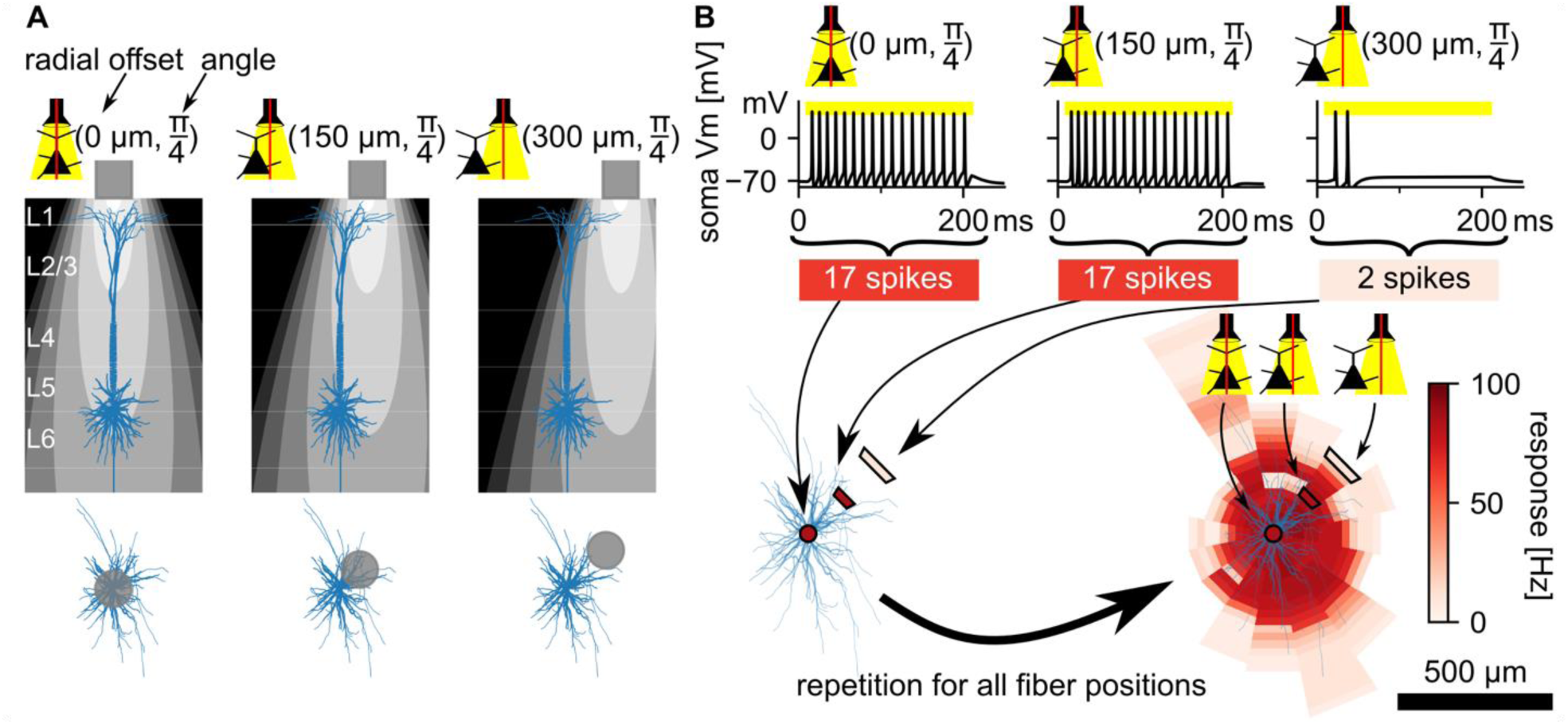
Heterogeneous distribution of stimulator locations induces strong optogenetic responses. **(A)** Illustration of the overlap of light intensity throughout cortical space with neuron morphology for three optical fiber locations increasing in radial offset from center. **(B)** Optogenetic responses to a 200 ms-duration stimulation at the example optical fiber locations shown in (A). The membrane voltage is measured at soma and thresholded at 0 mV to count elicited spikes. Spike counts are encoded with color and mapped spatially at the respective stimulator position to reveal the impact of neuron morphology (blue) on the dependence of optogenetic response on stimulator location. Repetition of the optogenetic response simulation at various optical fiber locations reveals the full spatial response profile. Stimulation intensity was 16 mW/mm² at the stimulator surface. Stimulator was an optical fiber with diameter of 200 µm and numerical aperture of 0.22.

### Increasing stimulation intensity shifts the origin of strongest ChR conductance

Next, we investigated the impact of varying stimulation intensity on the optogenetic responses of the layer-5 neuron. We varied the stimulator position along the cortical surface while stimulating at low, intermediate, and high stimulation intensities [Fig. 4A]. Recording the ChR conductance across all compartments, we found that the origin of strongest ChR conductance shifted with increasing stimulation intensity from the apical tuft to the apical shaft [Fig. 4B]. Contributions of single basal-dendritic, somatic, or axonal compartments to total ChR conductance were negligible. Reasons for the shift were two-fold. First, ChR channels in the apical tuft saturated with increasing light intensity earlier than in the apical shaft as apical tuft compartments were exposed to comparably higher light intensity being located closer to the stimulator. Second, a higher surface area in the apical shaft accommodated a higher number of channels, which could in total sum up to a larger ChR conductance per compartment although being exposed to less light. See Supplementary Materials Figure S1 for more details. The shift of strongest ChR conductance (per compartment) from apical tuft at low to the apical shaft at higher stimulation intensity resulted in a change of the spatial distribution of the neuron’s spiking response [Fig. 4C]. At low stimulation intensity (1.6 mW/mm²), spiking responses appeared only if the stimulator was located where the apical tuft reached close to the cortical surface, i.e., in isolated and off-center stimulator locations [Fig. 4C, left]. At higher light intensities (16 mW/mm² and more) for which the apical shaft contributed the strongest ChR conductance per compartment [Fig. 4B middle], spiking responses were highest for central stimulator positions, for which the apical shaft received the highest light exposure [Fig. 4C, middle]. These results suggest that for spatial precision, stimulation of layer-5 pyramidal neurons with a wide branching apical tuft is unfavorable at low stimulation intensities, as neurons located laterally to the light source might receive stronger optogenetic excitation compared to neurons located right below. Increasing the stimulation intensity could mitigate this effect, as it recruits more ChR conductance in the apical shaft, which resulted in an optogenetic response spatially centered around the neuron’s soma in our simulations.

**Figure 4:**
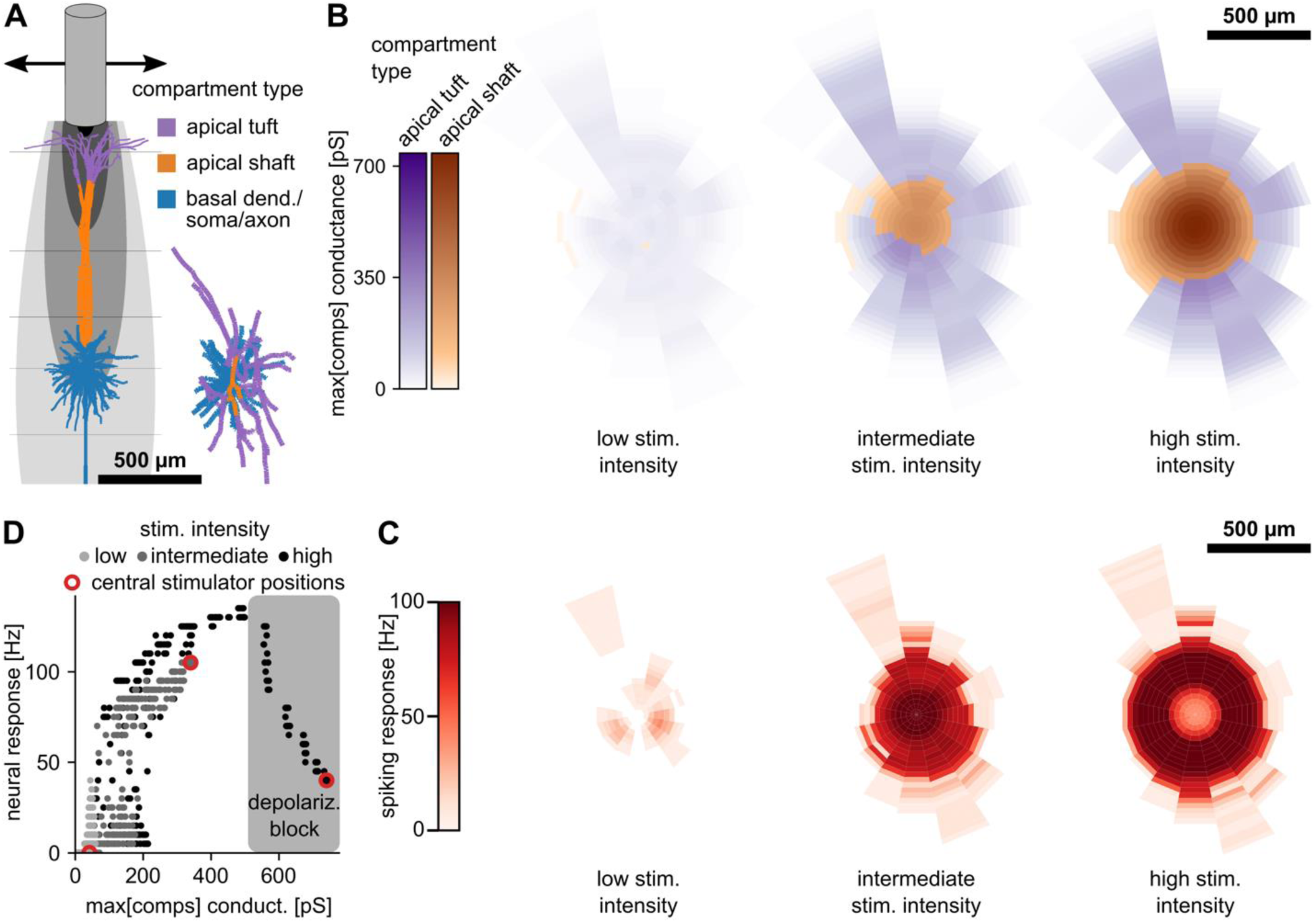
Optogenetic responses redistribute with increasing stimulation intensity. **(A)** Apical tuft (purple), apical shaft (orange), and remaining neuron morphology (blue) of the layer-5 pyramidal neuron and stimulator light profile inside the cortex viewed from the side (left) and neuron morphology viewed from the top (lower right). **(B)** Dependence of maximum channelrhodopsin conductance in a single compartment on stimulator location for low (1.6 mW/mm²), intermediate (32 mW/mm²), and high stimulation intensity (160 mW/mm²). Color encodes compartment type where maximum conductance is reached. Basal dendritic, somatic, and axonal compartments had negligible contributions and therefore were not displayed for visualization purposes. **(C)** Optogenetic spiking response depending on stimulator location at low, intermediate, and high stimulation intensity. **(D)** Strongest ChR conductance observed across compartments for varying stimulator locations (central location marked by red circle, otherwise not encoded) and stimulation intensity (color-coded). Depolarization block is induced only for the highest stimulation intensity condition and is strongest in the central stimulator location.

### Depolarization block preferentially occurs at central stimulator locations causing a ring-shaped response

At high stimulation intensity, depolarization block additionally reshaped the spatial characteristics of the optogenetic response [Fig. 4B,C,D; Supplementary Materials Fig. S2]. Central stimulator locations, which recruited the strongest ChR conductance per compartment, induced depolarization block with increasing stimulation intensity before other locations. As recruited maximal ChR conductance decreased with stimulator offset from center, depolarization block was induced inside a ring around the neuron’s soma, which increased in radius with stimulation intensity. These observations indicate that neurons below the optogenetic stimulator can be suppressed due to depolarization block while neurons located laterally to the stimulator still respond with strong sustained responses at high stimulation intensities.

### Stimulation recruits neurons in layer 5 and layer 2/3 differently

Since our simulations indicated that morphology causes complex optogenetic responses, we were interested whether this implies distinct layer-specific optogenetic activation patterns, as a neuron’s morphological type is linked to its layer-origin. We therefore simulated the stimulator-location-dependent optogenetic responses of a layer-2/3 pyramidal neuron to compare them to the layer-5 neuron’s responses [Fig. 5A]. In contrast to the layer-5 neuron, optogenetic response maxima were evoked for centered stimulator locations also at low and not only at intermediate stimulation intensities. Depolarization block was, as in the layer-5 neuron, first induced at centered stimulator locations when stimulation intensity was increased. However, the thresholds for response and depolarization block were at different stimulation intensities compared to the layer-5 neuron. Consequently, the spatial response characteristics of the layer-2/3 and layer-5 neurons differed when stimulated at the same stimulation intensity [Fig. 5A]. We found inverse response dependencies on radial stimulator offset at low stimulation intensity (3.2 mW/mm²) as response maxima were evoked by off-center stimulator locations in the layer-5 while evoked by central stimulator locations in the layer-2/3 neuron. At increased stimulation intensity (9.6 mW/mm²), response maxima were evoked at central stimulator positions for both neuron types. A further increase (32 mW/mm²) induced depolarization block at central stimulator positions only in the layer-2/3 neuron, while the layer-5 neuron retained a sustained response. At high stimulation intensity (160 mW/mm²), depolarization block occurred in both neurons for central stimulator locations, however, affecting the layer-2/3 neuron within a wider area of stimulator locations. These observations indicate that differences in morphology due to layer-origin of a neuron give rise to a spatially different distribution of optogenetic excitation in populations of layer-5 and layer-2/3 neurons, which we visualized accounting for different neuronal densities of these neurons in cat visual cortex (27) in Fig. 5B. In summary, our results demonstrate that stimulation intensity not only controls response strength but also modulates spatial response distribution [Fig. 5C].

**Figure 5:**
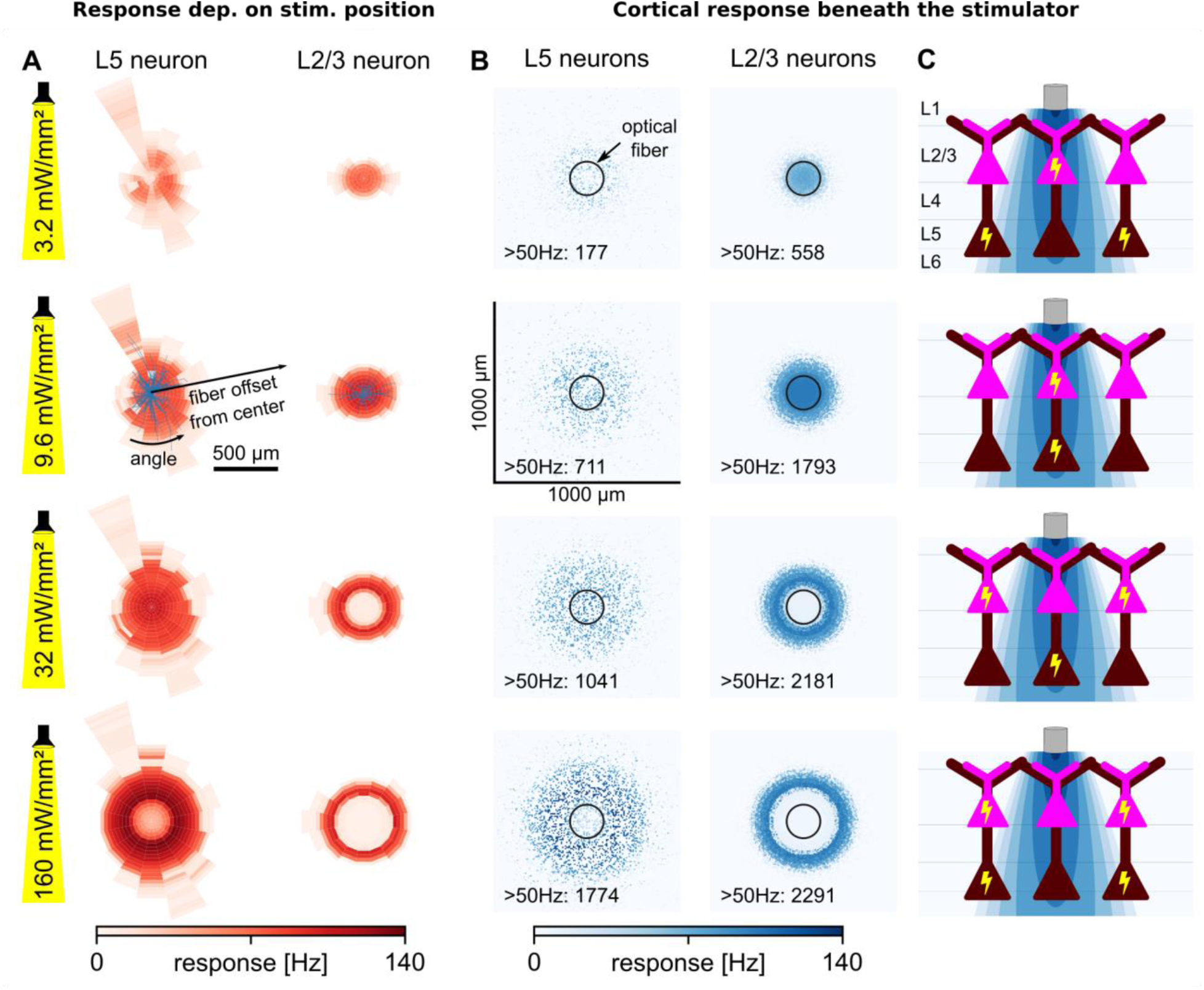
Spatial distribution of optogenetic excitation is dependent on the layer-origin of the neuron. **(A)** Optogenetic response dependence on stimulator location for layer-5 (left) and layer-2/3 pyramidal neuron (right) at varying stimulation intensity (rows). **(B)** Activated layer-5 (left) and layer-2/3 pyramidal cells inside 1 mm² of cortex around a stimulator (optical fiber) located in the center (black circle) and accounting for neuron-type-dependent density (27). **(C)** Qualitative illustrations of the different spatial distributions of optogenetic excitation in layer-5 (brown) and layer-2/3 (pink) neurons beneath the stimulator and depending on stimulation intensity (rows).

### Quantification of neuron-type-specific spatial stimulation precision

Since optogenetic responses varied in their spatial extent with varying stimulation intensity, and therefore response strength, we quantified stimulation precision in layer-5 vs. layer-2/3 neurons dependent on their activation level. [Fig. 6]. We varied stimulation intensity starting from below response-threshold all the way to a level that exceeded the threshold for depolarization block. For each simulated stimulation intensity step, we first averaged the optogenetically induced responses across the angular stimulator coordinate, and then found the maximum response in the resulting distance-dependent profile, which we termed “peak response” and used as indicator for the neuron’s response strength. Next, we calculated the stimulator distance at which the neuron’s response dropped to half of the peak response, which we defined as “response space constant” and utilized as a measure of stimulation precision [Fig. 6A].

**Figure 6:**
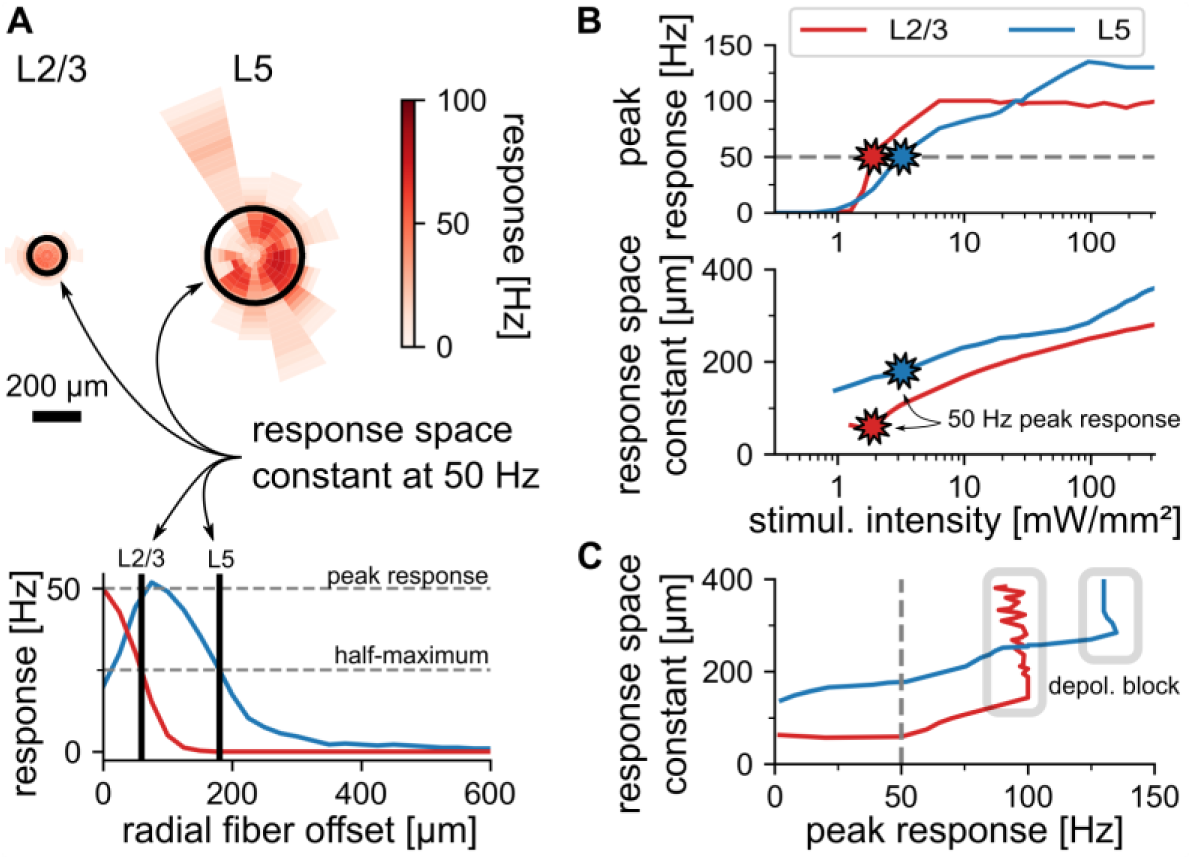
Comparison of spatial stimulation precision of layer 5 and 2/3 neurons at equal target response levels. **(A)** Examples of optogenetic response profiles and response space constant at a target response level of 50 Hz (top). Determination of “peak response” (maximum) and “response space constant” (radial fiber offset at half-maximum) from the stimulator-offset dependent optogenetic response. **(B)** Peak response of layer-2/3 (red) and layer-5 neuron (blue) plotted over stimulation intensity at the stimulator output (top). Response space constant for both neurons plotted over stimulation intensity (bottom). **(C)** Response space constant plotted against targeted peak response, same color-coding as in (B). The response space constant is not directly dependent on peak response, instead both variables are dependent on the underlying variable stimulation intensity. Diverging of the response space constant at certain peak response level is occurring due to depolarization block at high underlying stimulation intensities.

The peak response increased monotonically with stimulation intensity in both neuron types [Fig. 6B, top]. However, layer-5 and layer-2/3 neuron had different response thresholds, gains, and saturation points, which meant that at lower stimulation intensity, the layer-2/3 cell responded with higher firing rate than the layer-5 cell and vice versa at higher intensities. Crucially, the response space constant increased with stimulation intensity in both neuron types [Fig. 6B, bottom]. Hence, higher stimulation intensity resulted in a stronger response but also decreased spatial precision. Finally, we plotted the response space constants over the peak response, to provide a comparison of stimulation precision at equal response levels for the two examined neuronal types [Fig. 6C]. We found that throughout the response range below depolarization block, layer-2/3 neurons can only be engaged above half-maximum within about 100 µm distance between stimulator and soma while layer-5 neurons can be engaged within about 200 µm to this strength.

### Restricting ChR expression to the soma improves spatial stimulation precision only in the layer-5 neuron

An intuitive measure to prevent the activation of neurons from distant stimulator positions due to illumination of their dendritic arbors is restricting the expression of optogenetic constructs to the cell body. Shemesh et al. (26) have experimentally demonstrated that an adapted version of the CoChR protein, which selectively traffics to the cell body of neurons, improves the spatial stimulation precision of holographic optogenetic stimulation in cell cultures and cortical slices. To understand if such soma-targeting of the opsin is likely to improve the precision when the stimulation is applied with a divergent light source from the cortical surface, we replicated their measured soma-targeted opsin expression distribution in addition to the default uniform distribution representing no targeting in our simulations [Fig. 7A,B; see Methods]. We simulated two alternative somatic targeting conditions as their data were not clear on the presence of residual ChR transfection in the distal dendrites. First, we fitted an exponential decay plus y-offset plateauing at about 20% of the density expressed at the soma (“soft soma targeting”), and second, an exponential and linear decay plus y-offset which decays to zero density at around 200 µm distance from soma (“strict soma targeting”). In addition, we explored the impact of different absolute ChR expression levels in addition to different relative ChR distributions. The default ChR expression density used in our simulations was 130 channels/µm², which matched an empirical estimate of bacteriorhodopsin density in *Xenopus oocytes* (28). We additionally tested a 10-fold higher expression density (1300 channels/µm²).

**Figure 7:**
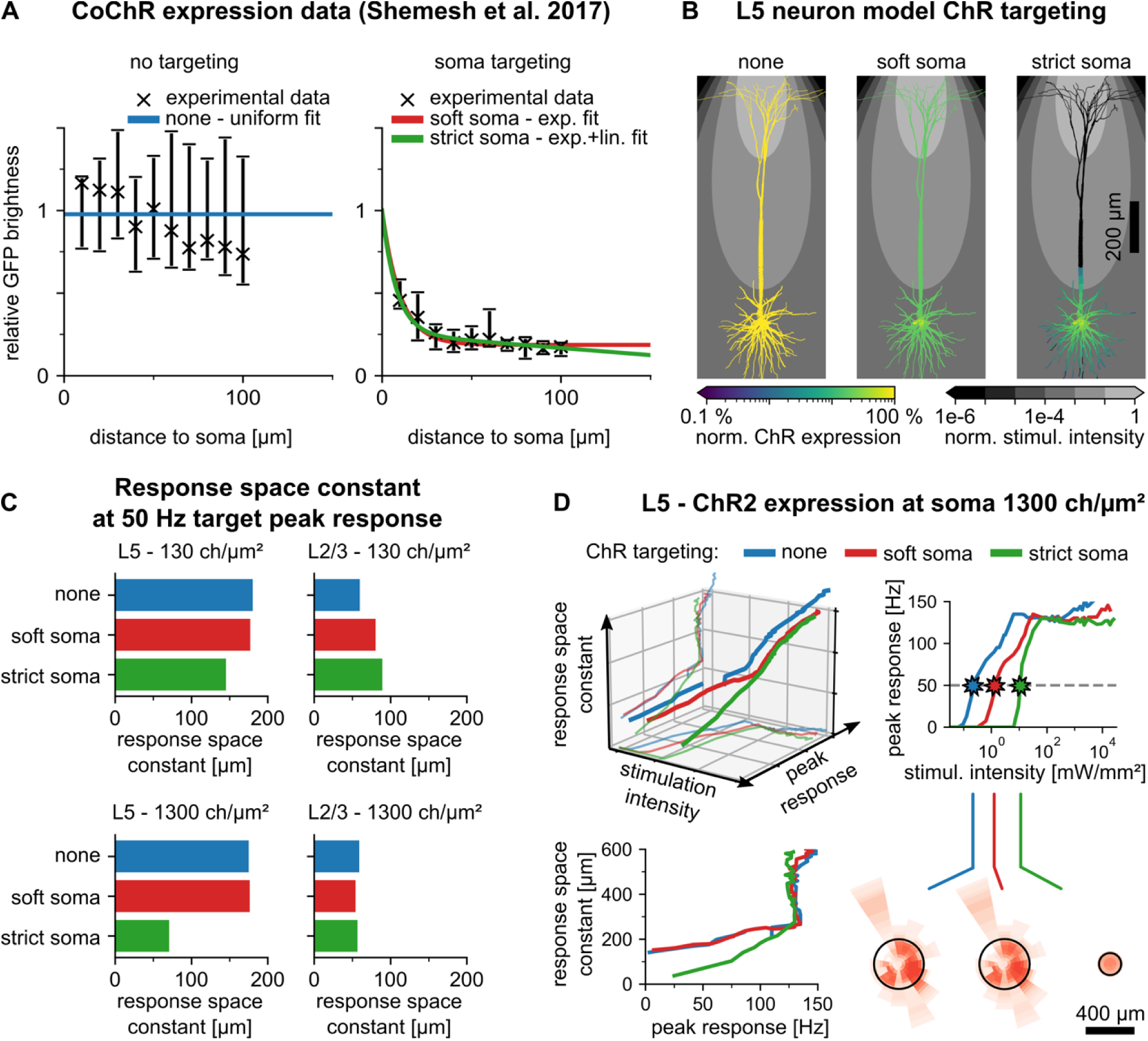
Preferential somatic ChR expression improves spatial precision only in the layer-5 pyramidal neuron. **(A)** Normalized channelrhodopsin expression distributions (solid lines) fitted to GFP-fluorescence measurements (discrete data, mean and std) (26). **(B)** Expression distributions from (A) applied to the layer-5 pyramidal neuron. **(C)** Comparison of response space constants observed at 50 Hz target peak response for the layer-5 and layer-2/3 pyramidal neuron type at channelrhodopsin expression levels of 130 channels/µm² and 1300 channels/µm². **(D)** Detailed results of the effect of somatic targeting for the layer-5 neuron and an expression level of 1300 channels/µm² for which strict somatic targeting achieved the strongest improvements in spatial precision. The 3D plot (upper left sub-panel) illustrates the dependence of peak response and response space constant on stimulation intensity. The bottom projection shows the dependence of the peak response on stimulation intensity, which is also plotted in 2D in the upper right sub-panel. Stars in the upper right sub-panel mark the data points at 50 Hz target peak response, from which the bar plot in (C) was generated. Spatial response profiles corresponding to these data points are plotted below (lower right sub-panel). The lower left sub-panel shows the behavior of the response space constant with peak response, which is also plotted as projection in the 3D plot (upper left sub-panel).

We compared spatial precision between conditions by comparing the response space constant at the same target peak response level. Somatic targeting improved spatial precision only for the layer-5 neuron, and required strict somatic targeting of ChR, as shown for an example target response level of 50 Hz in Figure 7C. Higher ChR expression (1300 channels/µm²) further enhanced the precision. As notable precision improvement was restricted to the layer-5 neuron with strict somatic targeting of ChR at high expression density (1300 channels/µm²), we included detailed simulation results only for this condition in Figure 7D. Detailed results for the other conditions can be found in Supplementary Materials Fig. S3. Across conditions (neurons, targeting, ChR expression level), somatic targeting required an increase of stimulation intensity, which amounted to two orders of magnitude in the layer-5 neuron and one order of magnitude in the layer-2/3 neuron [Fig. 7D, Supplementary Materials Fig. S3]. Symmetry of the stimulator-location-dependent response profiles was improved with strict somatic targeting in both neurons [Fig. 7D, Supplementary Materials Fig. S3]. In conclusion, our results suggest that somatic targeting of ChR can improve spatial precision only in the layer-5 neuron, but symmetry in both neurons, at the cost of a strong increase of light irradiance. Its effect further requires zero residual ChR expression in distal dendrites.

To understand whether the reported effects of ChR targeting generalized across optical fiber parameterizations, we repeated the simulations for the “no targeting” and “strict targeting” conditions and evaluated the response space constant at a peak response of 50 Hz. Indeed, the substantial improvement of spatial precision through strict targeting of ChR in the layer-5 neuron and the lack of improvement in the layer-2/3 neuron were present for most simulated optical fiber parameterizations (fiber diameters d = [25, 400] µm, numerical apertures NA = [0.1, 0.5]) [Fig. 8].

**Figure 8:**
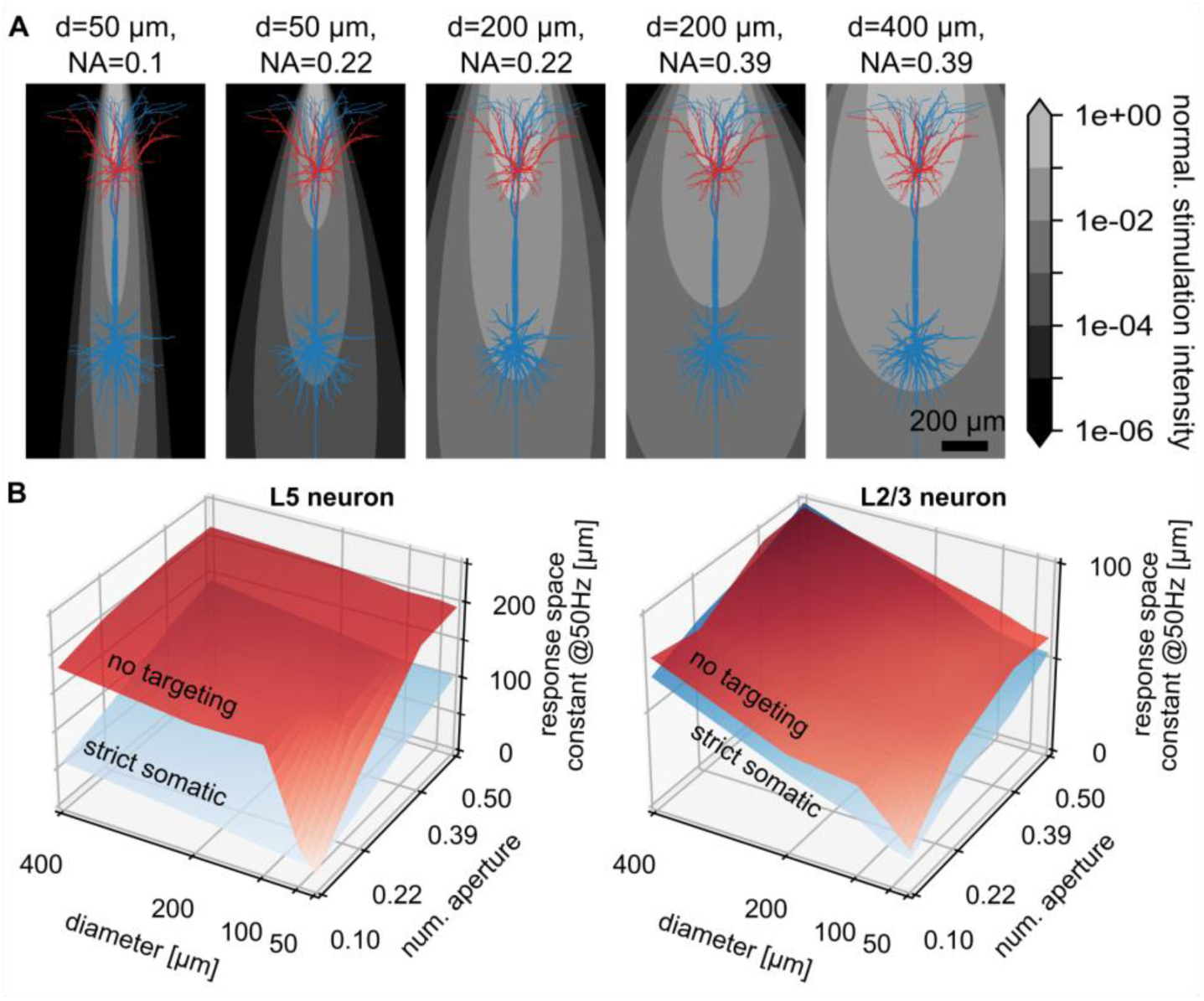
Narrowing optical fiber diameter and numerical aperture improves stimulation precision. **(A)** Light intensity levels throughout cortical space with layer-5 and layer-2/3 pyramidal neuron superimposed for several example optical fiber parameterizations. **(B)** Dependence of the response space constant at a target peak response of 50 Hz on optical fiber diameter and numerical aperture for the layer-5 and layer-2/3 neuron and using no (red) and strict somatic targeting of ChR2 (blue).

### Narrowing the stimulator light profile improves precision independently of neuron-type

Narrowing the stimulator’s light profile decreased the spatial extent of the optogenetic response in both neuron types [Fig. 8]. The relative improvement of their response space constants between the widest (d=400 µm, NA=0.5) and narrowest (d=25 µm, NA=0.1) optical fiber conditions was approximately four-fold for the layer-2/3 neuron and three- or four-fold for the layer-5 neuron, depending on whether somatic targeting was involved or not. With the optical fiber narrowed to the tightest light beam profile explored (d=25 µm, NA=0.1), stimulation precision reached its maximum regardless of whether somatic targeting of channelrhodopsin was involved or not.

## Discussion

Cortical implants utilizing optogenetic stimulation to engage functionally meaningful neuronal populations are posed to play a key role in unraveling of how cortical circuits process information (5–7, 11, 12), how functional network architecture develops (29, 30), and in an applied form of neuro-prosthetic devices for restoration of sensory impairments, e.g., vision (9, 10, 15). Success of these endeavors will require optogenetic stimulation being precise at a scale of at least hundreds of micrometers, at which many functional neuronal representations are encoded (e.g., orientation preference in primary visual cortex of higher mammals (13)). We demonstrated here with computer models which are supported by experimentally observed response dynamics that shape, size, and cortical depth of neuronal morphology impact stimulation precision. Importantly, we found that these impacts depend on stimulation intensity in a highly non-linear fashion, as increased stimulation intensity could result in both elevated and decreased neuronal response depending on the exact intensity, which has several practical implications for optogenetic experiments. We also used the model to understand how subcellular targeting of channelrhodopsin and stimulator optics can affect precision, finding general improvement across neuron types only through optical changes. Therefore, we lay foundations for interpreting previous and future experimental results as well as for designing future optogenetics-based implants in a more informed manner.

According to our simulations, optogenetic stimulation causes direct neuronal activation over a spatial width of several hundred micrometers. Specific predictions are significant stimulation of a layer-5 neuron at about 250 µm and a layer-2/3 neuron at about 100 µm of lateral distance between the stimulator and the neuron’s soma. These results are compatible with experiments in the primary visual cortex of tree shrew, which reported significant optogenetic activation within 150 µm and no activation beyond 300 µm when excitatory synaptic blockers were applied, and hence, only direct neuronal stimulation was permitted (5).

Our results have several implications, both for spatially precise application of optogenetic stimulation, and for the interpretation of optogenetically induced neuronal responses. First, stimulation at high light intensity caused optogenetic excitation to be suppressed by depolarization block centrally below the optogenetic stimulator while excitation remained strong in the surrounding of the suppressed region [Fig. 4]. Such specific unwanted geometry of activation can have unexpected effects that can easily lead to misinterpretation of the experimental results, while being difficult to detect. Predicting the light intensity at which depolarization block occurs might be challenging due to fluctuating levels of transfection, both between cells and between experiments. Hence, when increasing stimulation intensity to recruit neurons in a large volume, activation level in the center of stimulation should be monitored to detect and avoid suppression effects due to depolarization block. Alternatively, the appearance of depolarization block may be avoided through pulsed instead of continuous stimulation (31). According to our experimental and model results, depolarization block can appear at about 10 ms after stimulation onset, hence requiring pulsed stimulation at frequencies of 100 Hz or more to avoid its onset. Finally, depolarization block also offers an interesting opportunity to suppress neurons below the optogenetic stimulator on purpose while activating them on a ring around the suppressed location.

The second implication concerns the spatial distribution of optogenetic excitation, which we found varies with stimulation intensity in pyramidal neurons of layer-5 and layer-2/3 [Fig. 5A]. Therefore, modulation of stimulation intensity is not only affecting the neuronal response strength but can also serve as an instrument to optimize the spatial distribution of optogenetic stimulation [Fig. 5B,C]. However, stimulation of both neuron populations such that they responded centrally below the stimulator required restricting the stimulation intensity to a narrow range, which left little flexibility with neuronal target response levels.

Third, we found that not only the response level but also the distance at which a neuron is still stimulated increases with light stimulation intensity. Similarly, optogenetic excitation of inhibitory neurons in the cortex of mice yielded a concomitant increase of the inhibitory neurons’ response strength and spatial response extent with stimulation intensity (14). Since, synaptic activity was not blocked in these experiments, and the observed spatial response extent (> 1 mm) was larger than the one we found for stimulation of isolated single neurons, the observations taken together suggest that network mechanisms may amplify the effect being already present at single neuron level. Therefore, increasing the stimulation intensity to amplify neuronal responses comes at the cost of spatially broader neuronal responses. Vice versa, lowering the stimulation intensity could yield a spatially more specific neuronal response.

Fourth, we observed that somatic targeting of ChR can more than double the spatial stimulation precision in the layer-5 pyramidal cell, while it has negligible effects on stimulation precision in the layer-2/3 neuron. Crucially, strong improvement was present only at high expression of ChR (1300 channels/mm²) with no residual expression of ChR in the apical tuft, highlighting that improving precision only works with a potently expressing and effectively targeting optogenetic construct. Somatic targeting also required up to two orders of magnitude higher light intensities to reach the same target response level to compensate for the absence of ChR expressed in dendrites [Fig. 7B,C]. Such large increase of stimulation intensity may cause phototoxicity-mediated cell death in superficial layers. Similarly, the concern of phototoxicity applies to the improvement of precision by narrowing the stimulator light emission profile, which also relies on increasing light intensity to compensate for the decrease in stimulator output surface [Fig. 8]. However, concerns of phototoxicity may be alleviated through using opsins with higher light sensitivity, which allow for effective stimulation at lower stimulation intensities (32).

Various computational research investigated optogenetic stimulation of single neurons (18, 19, 33–35). However, to the best of our knowledge, only (18) and (19) have studied the spatial dependence of optogenetic stimulation on the relative position of the light source to the neuron. Specifically, these studies focused on characterizing the stimulation intensity threshold required to induce neuronal spiking. Their results highlighted that dendrites contribute to optogenetic activation and that altering the spatial distribution of ChR throughout the cell can lower the stimulation threshold. Our results are complimentary to theirs as we report on the spatial dependency of optogenetic responses in the regime beyond response threshold, which their studies omitted by design. Knowledge of this dependency is crucial for the design of spatially coordinated optogenetic interventions as it reveals how strongly a neuron is activated given its distance to the stimulator and role in the cortex (e.g., layer-2/3 vs. layer-5). However, covering only the direct activation through optogenetic stimulation, our work allows only for predictions on the effective optogenetic input to the neuronal network. This input is further reshaped by network-level effects, which necessitate detailed spiking network simulations to be understood. On one hand, previous simulations relying on the point-neuron model approximation found a spatial spread of activation extending beyond the spatial extent of the optogenetic stimulus for random connectivity (16). On the other hand, our previous simulation study of V1 cortex found an effective functional sharpening of the optogenetically delivered external input in the orientation domain due to a competitive network-level mechanism implemented in the cortical lateral connectivity (15). Future work needs to combine the reported single neuron response characteristics from this study with neuronal network modeling to enable a comprehensive exploration of the mechanisms underlying cortical optogenetic responses.

There are several limitations to the present study that warrant further research. Our model predicts the occurrence of depolarization block for high stimulation intensities matching experimentally observed dynamics only in about half of the experimentally recorded ganglion cells originating from one Macaque retina. The absence of depolarization block in the remaining half of the cells may be attributed to a lower opsin expression density, which could unfortunately not be experimentally verified. We computationally confirmed that lower opsin expression levels indeed require higher stimulation intensities to induce depolarization block [see Supplementary Materials Fig. S4]. However, further experimental evidence is needed to explore whether depolarization block appears more universally for high stimulation intensity across various neuron types.

To reach a complete understanding of how large neuronal populations respond to optogenetic stimulation, neurons with various other morphologies and biophysical properties need to be characterized in addition to the here characterized excitatory neurons from layers 2/3 and 5. Particularly, excitatory and inhibitory neurons may exhibit systematic differences between their optogenetic response characteristics, which may enable a richer repertoire of stimulation strategies given their functional different impact on network activation. Furthermore, morphology and biophysics vary among neurons of the same type. Our analysis could in future be applied to quantify these inter-neuron variations using computational models representing a larger variety of the same type of neuron. Application of our analysis to a wider set of similar neuron models would further help detecting potential errors in our models’ parameterizations, which although being well established, relied partly on only a small number of experimentally challenging measurements.

Another limitation of our study is that we did not account for synaptic background activity, which depending on its type (excitatory/inhibitory) could add up with or diminish charge introduced into the cell through photoactivation of ChR. On average this may result in a shift of the activation threshold but also, if the synaptic input is spatially biased, affect the spatial response characteristics. Furthermore, the current study only considers temporally steady light stimulation. Pulsed stimulation is a commonly used temporal protocol (see (33) for a comparison), thus studying how parameters of such protocol impact the response characteristics is of great interest. Another limitation of this work is that neuron models did not include axonal arbors which could significantly contribute to optogenetic response according to an earlier modeling study (18). Especially large axon-bundles, which extend over wide distances in horizontal direction as for example observed in layer 2/3 of the primary visual cortex of cats (36), could give rise to additional distant clusters of light source positions stimulating the neuron substantially. However, such response clusters have not been observed in experiments applying optogenetic stimulation from the cortical surface to the primary visual cortex of tree shrews (5). Future modeling work based on neuronal reconstructions including at least partial axonal arbors, could help clarify their contribution to optogenetic activation.

## Methods

### Animals

All experiments were conducted in accordance with the National Institutes of Health Guide for the Care and Use of Laboratory Animals. The protocol was approved by the local animal ethics committees and was conducted in accordance with Directive 2010/63/EU of the European Parliament. Cynomolgus macaques (Macaca fascicularis) of foreign origin were used for the study.

### Cell-attached recordings of primate RGCs

Experimental data on cell-attached recordings of primate retinal ganglion cells have been partially published previously with a detailed description of methodology (37).

#### AAV production and eye injection

AAV backbone plasmid contained human-codon-optimized CatCh sequence. SNCG promoter was cloned into the AAV backbone plasmid. The constructs further contained woodchuck hepatitis virus post-transcriptional regulatory element (WPRE) and bovine growth hormone poly (A).

Animals were anesthetized with ketamine/xylazine (10:1 mg/kg). The vitreous was injected with 100 µL of viral vector holding 10¹² viral particles. After injection, the cornea was treated with an ophthalmic steroid and antibiotic ointment.

#### Electrophysiology

AAV-treated macaque retinas were recorded in oxygenized (95% O2, 5% CO2) Ames medium (Sigma-Aldrich), to which a selective group III metabotropic glutamate receptor agonist, L-(+)-2-amino-4-phosphonobutyric acid (L-AP4, 50 mM, Tocris Bioscience, Bristol, UK) was added.

Cell-attached recordings were conducted with an Axon Multiclamp 700B amplifier, using borosilicate glass capillaries (BF100-50-10; Sutter Instruments), pulled to 7-9 MΩ. Retinal ganglion cells were recorded in current-clamp configuration (current zero) with electrodes filled with Ames’ solution. Retinal ganglion cells were dark-adapted for one hour prior to the recording.

Full-field photostimulation was performed with a Polychrome V monochromator (Olympus, Hamburg, Germany), and output light intensities and wavelengths were calibrated.

### Simulations and computational model

#### Simulation software

We simulated neuronal dynamics with the NEURON simulator, version 8.0.0 (20), through its python-interface (38) using python 3.6. We used the snakemake workflow manager (39) to organize the simulation workflow, to parallelize, and to manage runs with various parameter sets.

#### Neuron models

We simulated neurons with active dendrites using existing multi-compartment models based on reconstructions of a layer-5 and layer-2/3 pyramidal cell (21) and electrophysiological dynamics previously identified based on experimental recordings for the layer-5 cell (22). Somato-dendritic electrophysiological dynamics included voltage-gated sodium (na) and potassium channels (kv), slow non-inactivating (km), and calcium-dependent potassium channels (kca), as well as high-voltage activated calcium channels (ca). Axonal electrophysiology included low- and high-threshold sodium channels (na12/na16) as well as voltage-gated potassium channels (kv). Channels are implemented in the corresponding NMODL files (channel abbreviation + suffix ‘.mod’). To model the conductance through Channelrhodopsin-2 channels, we used a NMODL implementation of the channel by (18) (chanrhod.mod; for model details, see “Channelrhodopsin model”). As a default, we assumed a uniform density of 130 channels/µm^2^ throughout the neuron morphology (18, 28). This yielded 10354945 ChR2 channels for the layer-5 and 2588736 for the layer-2/3 cell (a fourth of the channels in the layer-5 cell). To investigate the effects of preferential expression of ChR2 at the soma (“soma targeting”), we set the somatic density to the otherwise uniformly applied density and decreased ChR density along the neuronal processes according to distributions derived from experimental data (cf. “Derivation of spatial channelrhodopsin expression distributions”). In addition to our default expression density of 130 channels/µm² we explored a ten-fold stronger expression (1300 channels/µm²) along with the simulations comparing somatic-targeting and no-targeting conditions (Figure 7, 8).

#### Channelrhodopsin model

The channelrhodopsin dynamics is based on a four-state Markov model describing the changes in conformation of the molecule (24, 40, 41) and has been previously implemented in NMODL (NEURON) (18). Model parameters are summarized in (18) and the original model publication (24). The model accounts for two open, i.e., ion-conducting, and two closed states. Under light illumination channelrhodopsin can transfer from the closed-1 to the open-1 state. From there it can transfer back to closed-1 or transfer to another open state, open-2, in which its conductance is lower than in open-1. From the open-2 state the molecule can transfer back to open-1 or to closed-2. The closed-2 state can thermally decay into closed-1 or under light illumination photoexcite back into open-2.

#### Optical fiber light model

We simulated the propagation of light emitted by an optical fiber inside the cortical tissue accounting for absorption and scattering of photons (absorption coefficient 0.125 mm⁻¹, scattering coefficient 7.37 mm⁻¹ (18, 42); index of refraction n=1.36 (43)) according to the Kubelka-Munk general theory of light propagation (18, 23, 43, 44). The model enables the simulation of various fiber models by adapting the parameters diameter and numerical aperture. The diameter defines the size of the output surface, and the numerical aperture defines the divergence of the emitted light beam. In our simulations we assumed that the optical fiber is placed on top of the cortical surface with its output surface directly connected to the cortical medium. We did not account for light losses involved in the coupling of the light into the cortical tissue. We used a fiber with a 200 µm diameter and numerical aperture of 0.22 as default and explored variations of the diameter of 25-400 µm and of the numerical aperture of 0.1-0.5 specifically [Fig. 8].

#### Derivation of spatial channelrhodopsin expression distributions

We derived the spatial distribution of ChR inside the neuronal morphology from experimental data obtained by (26). In this study, mouse cortical neurons were transfected with CoChR (Chloromonas oogama channelrhodopsin) with and without using somatic targeting motifs to restrict the channelrhodopsin expression to the soma. They attached green fluorescent protein (GFP) to CoChR to derive the relative surface density of CoChR through fluorescence measurements, which were taken in 10 µm steps along neurites and normalized to the fluorescence level at the soma.

In the condition without somatic targeting, the fluorescence measurements along the neurites had a slight decreasing trend with distance from soma. Considering their uncertainty, they were compatible with a uniform expression throughout the morphology, which we chose as distribution to model the “no targeting” condition [Fig 7 A, left]. A uniform distribution in the “no targeting” case is also supported by similar fluorescence measurements for a ChR2-YFP compound in cultured neurons from rats (25).

Somatic targeting of CoChR resulted in an exponential drop of the relative fluorescence with distance to the soma. However, it was not clear whether the exponential decay plateaued or further decayed beyond their most distant data point at 100 µm distance from soma. To account for these two alternatives, we fitted two distribution models to their soma-targeting data [Fig. 7 A, right]: Distribution model 1 represented a relative somatic targeting of ChR. The density decayed exponentially with the distance from soma until it plateaued at about 50 µm distance to soma at a level of about 20% of the somatic density. Since this model represented only an overexpression of ChR at the soma compared to distant dendrites, we referred to it as “soft soma targeting”. Distribution model 2 accounted for a complete decay of the ChR density. It described a similar exponential decay as distribution model 1 but had an additional linear decay which yielded zero ChR density at about 200 µm distance to the soma. We referred to this case as “strict soma targeting”.

#### Analysis of simulated optogenetic responses

To obtain spatial profiles of optogenetic responses exhibiting their dependence on the stimulator placement on top of the cortical surface, we simulated optogenetic stimulation with a constant intensity of 200 ms duration. We recorded the membrane voltage at the neuron’s soma, interpolated the data at 0.1 ms, and thresholded at 0 mV to count spikes. We converted the spike count to firing rate by dividing by the stimulation duration. We repeated this procedure at stimulator locations starting at the central location above the neuron’s soma and displacing the stimulator in lateral steps of 25 µm up to 975 µm. We additionally varied the stimulator location along angular direction in 16 steps along the full circle.

We decided to compare various conditions requiring the same target response level. We determined the response level as the maximum of the optogenetic response across the stimulator’s radial distance coordinate and averaged over its angular coordinate values. Since the response level is dependent on the stimulation intensity, finding a specifically targeted response level required a search algorithm. To find this target response level, we first mapped the optogenetic response for the layer-2/3 and layer-5 neurons on a logarithmic and coarse scale to determine the stimulation intensity thresholds for response and for depolarization block. Using these thresholds as limits, we used a bisection algorithm to find the stimulation intensity resulting in the desired target response level. The algorithm used as a first test value the center between the limits on a logarithmic scale and repeated this procedure while updating the upper/lower limit with the previously tested value depending on whether the resulting response level exceeded or fell below the targeted response level. During this procedure, the target response level was determined from the stimulator-distance-dependent optogenetic response profile only up to a radial distance of 200 µm to accelerate the search. We terminated the optimization procedure when the simulated response level was within 10% tolerance of the targeted one or terminated and discarded the data point if the target response level was not found within 31 steps.

Finally, we measured spatial precision of the stimulation by determining the radial stimulator distance at which the optogenetic response across radial stimulator distance and averaged over the stimulator’s angular coordinate reached half of the response level. We used linear interpolation if the stimulator distance yielding half of the response level was not covered by sampling.

## Acknowledgements and Funding Sources

We thank Thomas Foutz for technical advice on the multi-compartment neuron model and thank Lawrence Humphreys and Jose-David Celdran for helpful discussions on subcellular targeting of opsins.

## Authors contributions

Conceptualization, D.B. and J.A.; Methodology, D.B., L.B., A.C., G.G., S.P., and J.A.; Software, D.B., L.B.; Investigation, D.B. and A.C.; Writing – Original Draft, D.B. and J.A.; Writing – Review & Editing, D.B., L.B., A.C., G.G., and J.A.; Funding Acquisition, G.G., S.P., and J.A.; Resources, G.G., S.P., and J.A.; Supervision, G.G., S.P., and J.A..

## Data and materials availability

Electrophysiological data files are contained in Supplementary Materials. All original simulation code has been deposited at https://github.com/CSNG-MFF/osmorph/ and is publicly available as of the date of publication. Any additional information required to reanalyze the data reported in this paper is available from the corresponding authors upon request.

## List of Supplementary Materials

- Fig. S1: Mechanisms causing the shift of ChR conductance from the apical tuft to the apical shaft with increasing stimulation intensity.
- Fig. S2: Dependence of the optogenetic response pattern on stimulation intensity.
- Fig. S3: Detailed simulation results on the effect of somatic targeting on spatial optogenetic precision.
- Fig. S4: Lower opsin expression prevents occurrence of depolarization block.
- Data S5: Electrophysiological recordings of a Macaque retinal ganglion cell undergoing optogenetic stimulation at varying stimulation intensity visualized in Fig. 2 (RGC 1).
- Data S6: Electrophysiological recordings of a Macaque retinal ganglion cell undergoing optogenetic stimulation at varying stimulation intensity visualized in Fig. 2 (RGC 2).

**Figure S1:**
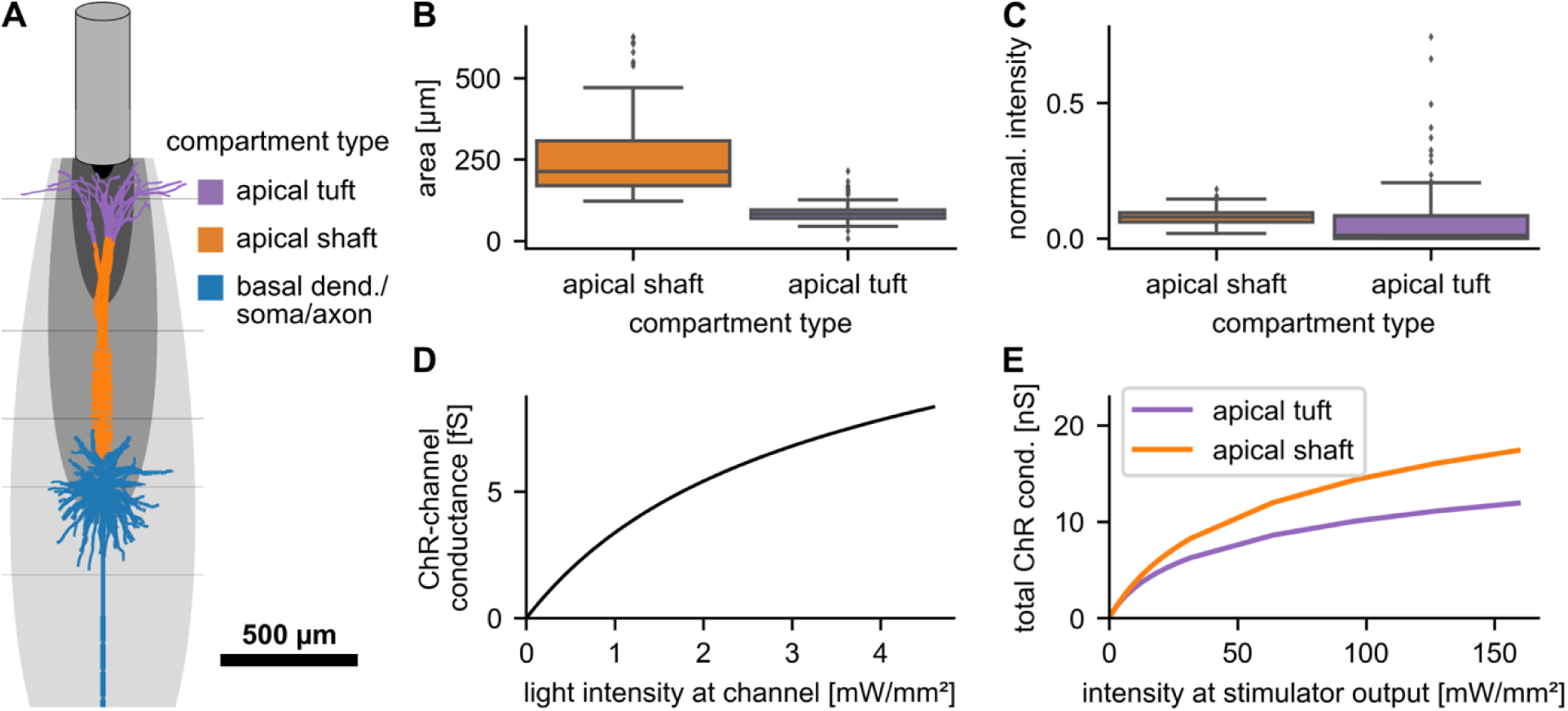
Mechanisms causing the shift of ChR conductance from the apical tuft to the apical shaft with increasing stimulation intensity. (A) Apical tuft (purple), apical shaft (orange), and remaining neuron morphology (blue) of the layer-5 pyramidal neuron and stimulator light profile inside the cortex viewed from the side. (B) Area per compartment tends to be larger in the apical shaft compared to the apical tuft. (C) Median light intensity is largest for apical shaft compartments, but strongest intensities are reached for apical tuft compartments. Values were calculated for the central stimulator position as an example. (D) Average conductance per ChR-channel increases sub-linearly with increasing light intensity. Therefore, the gain of ChR conductance decreases with increasing light intensity. Light intensity was here measured at the channel surface and conductance calculated as the temporal mean conductance within a 200-ms constant stimulus. (E) Total conductance through ChR increases more strongly in the apical shaft than in the apical tuft and therefore dominates the response at higher stimulation intensities.

**Figure S2:**
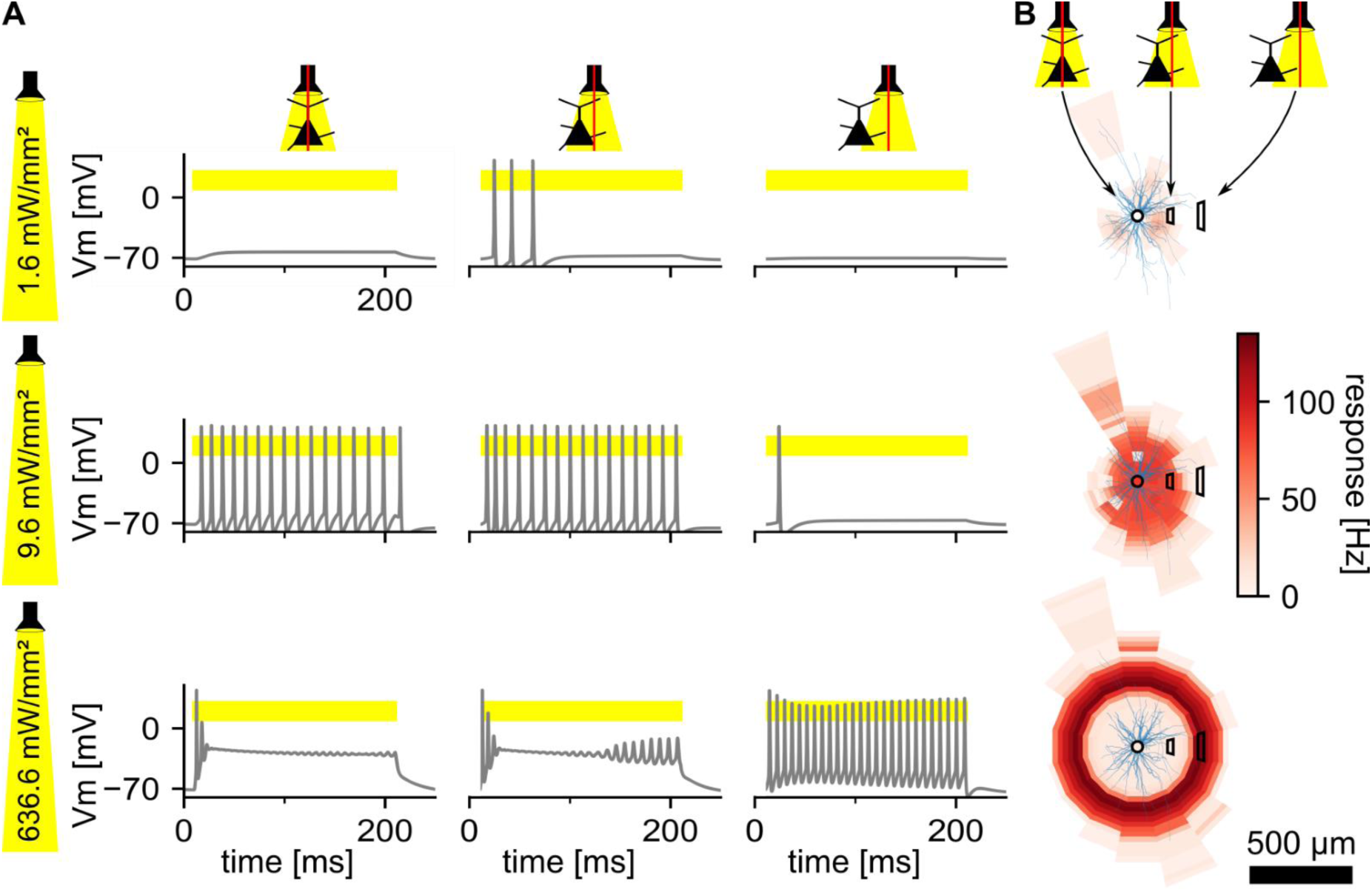
Dependence of the optogenetic response pattern on stimulation intensity. (A) Membrane voltage at the soma of the layer-5 neuron undergoing optogenetic stimulation (yellow) of varying intensity and location. Stimulator located on top of the cortical surface at radial distance of 0 µm, 150 µm, or 300 µm from soma (left to right). (B) Optogenetic responses depending on stimulator position. Three example stimulator locations from (A) marked with black frames and neuronal response encoded in color.

**Figure S3:**
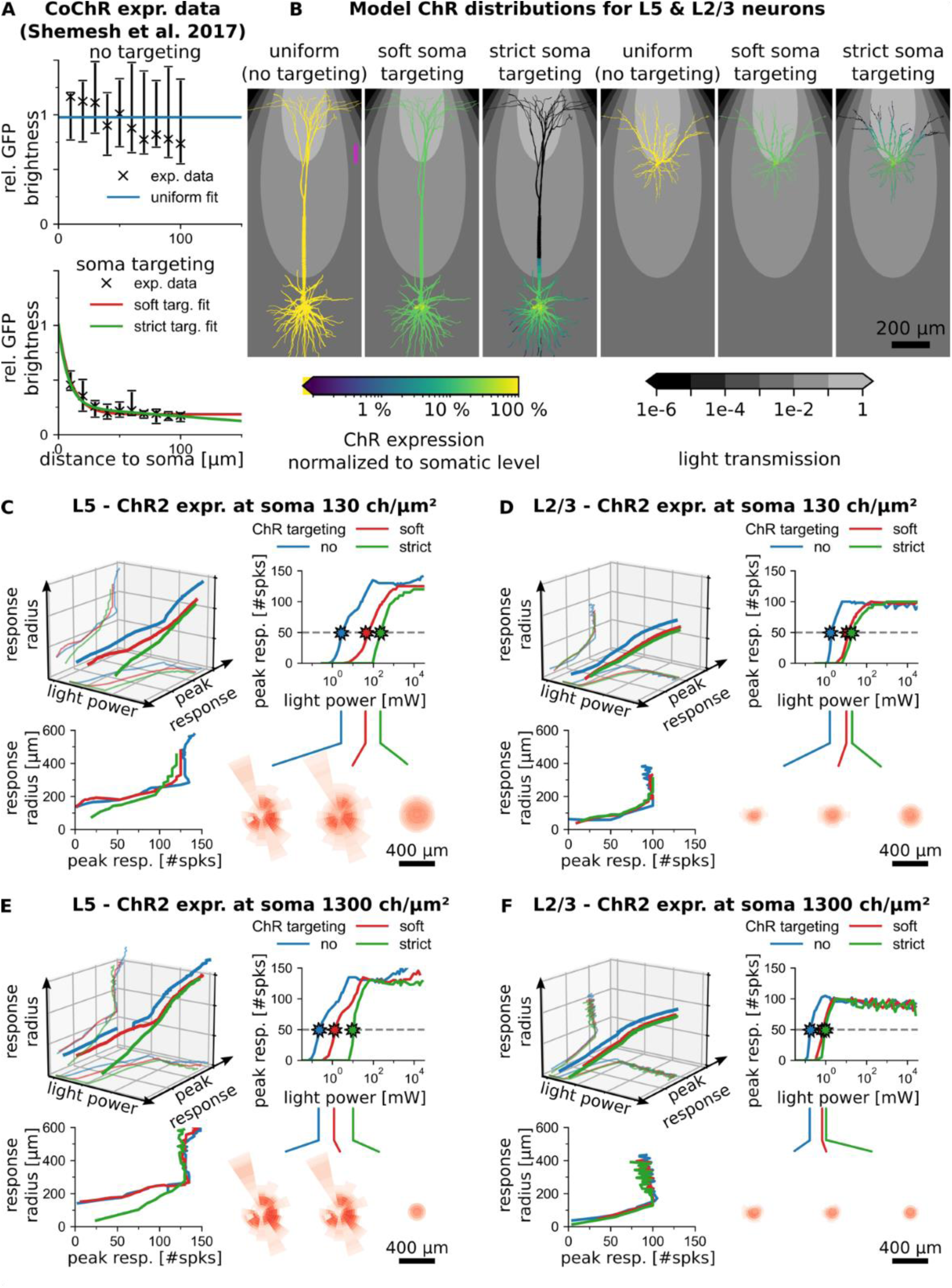
Detailed simulation results on the effect of somatic targeting on spatial optogenetic precision. (A) Normalized channelrhodopsin expression distributions derived from GFP-fluorescence measurements (23). (B) Expression distributions from (A) applied to the layer-5 and layer-2/3 pyramidal neuron. (C) Effect of somatic targeting for the layer-5 pyramidal neuron and a ChR expression density of 130 channels/µm² at soma. Behavior of the response space constant and peak response with stimulation intensity plotted in 3D (top left) with projections of peak response vs. stimulation intensity and response space constant vs. peak response plotted individually (top right, bottom left). The bottom right shows the stimulator-location-dependent optogenetic response for the three targeting conditions (no/soft/strict targeting) at a peak response of 50 Hz. (D) Same as (C) for the layer-2/3 pyramidal neuron. (E,F) Same as (C) and (D) at a 10-fold higher ChR expression density of 1300 channels/µm² at soma.

**Figure S4:**
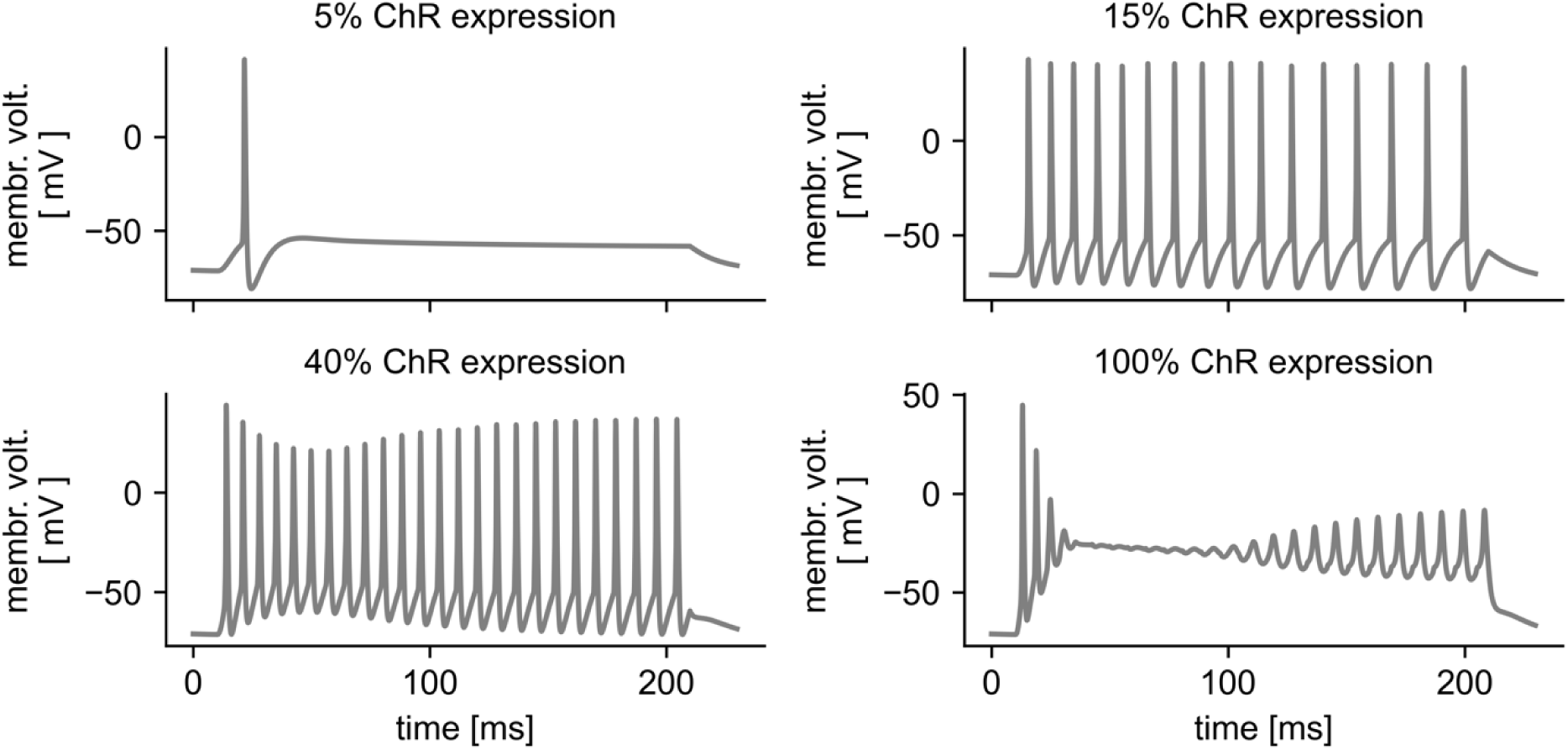
Lower opsin expression prevents occurrence of depolarization block. Panels show the voltage response of the layer-5 pyramidal neuron at the soma while the stimulation intensity was kept constant at 300 mW/mm² (at stimulator output surface) and ChR2 expression density was increased from left to right and top to bottom in approximately equidistant steps on a logarithmic scale. Stimulator diameter was 200 µm and numerical aperture 0.22.

